# Modified crRNA variants for precise targeting of point mutations in human mitochondrial DNA by CRISPR-Cas12a

**DOI:** 10.64898/2026.09.09.750411

**Authors:** Nikita Shebanov, Anne-Marie Heckel, Natalia Nikitchina, Anna Pavlova, Inna Pyshnaya, Mariya Meschaninova, Ivan Tarassov, Nina Entelis

**Author notes:** To whom correspondence should be addressed. N. Entelis; or I. Tarassov.

## Abstract

The CRISPR-Cas12a system, primarily used for nuclear genome editing, operates through RNA-guided cleavage of target DNA. Given the high copy number of mitochondrial DNA (mtDNA) in cells and the pathogenic potential of mtDNA mutations, there is substantial interest in developing methods to selectively eliminate mutant mtDNA without affecting wild-type molecules. We suggest using AsCas12a, with its shorter CRISPR RNA (crRNA) and proven ability to be targeted into human mitochondria, to perform selective cleavage of mutant mtDNA. Here, we assess the potential of AsCas12a and various engineered crRNAs to discriminate between wild-type and mutant mtDNA *in vitro* and in living human cells. By exploring chemically synthesized full and split crRNAs with various modifications, we identified crRNAs capable of targeting specific mutant mtDNA. We demonstrated the strong influence of magnesium concentration on the ability of crRNAs to discriminate point mutations. Finally, we show that AsCas12a-fused cytosine base editor can be addressed to mitochondria and can direct low but significant specific base editing of mtDNA in human cells. Our study suggests AsCas12a as a promising tool for targeted mutant mtDNA elimination or editing.

## INTRODUCTION

Human mitochondrial DNA (mtDNA) is a circular molecule of 16 569 bp which encodes 13 subunits of the oxidative phosphorylation complexes along with 22 tRNAs and 2 rRNAs required for their translation (**Fig. 1A**). There are hundreds to thousands of mtDNA molecules per mammalian cell, depending on the cell type. Mutations of mtDNA (mostly point mutations or deletions) can lead to mitochondrial dysfunction and cause severe neuromuscular diseases, whose manifestation and severity depend on the percentage of mutant molecules in the cell, so-called heteroplasmy level (1). Up to date, more than 350 pathogenic mtDNA mutations have been identified at the origin of various neuromuscular diseases (2). The simultaneous presence of mutant and wild-type mtDNA molecules opens the possibility of eliminating mutant molecules by targeting them with various enzymes, i.e. restriction enzymes (when mutations create a restriction enzyme site), Zinc-finger nucleases (ZFNs) or Transcription activator-like effector nuclease (TALENs) (reviewed in (3)).

**Figure 1:**
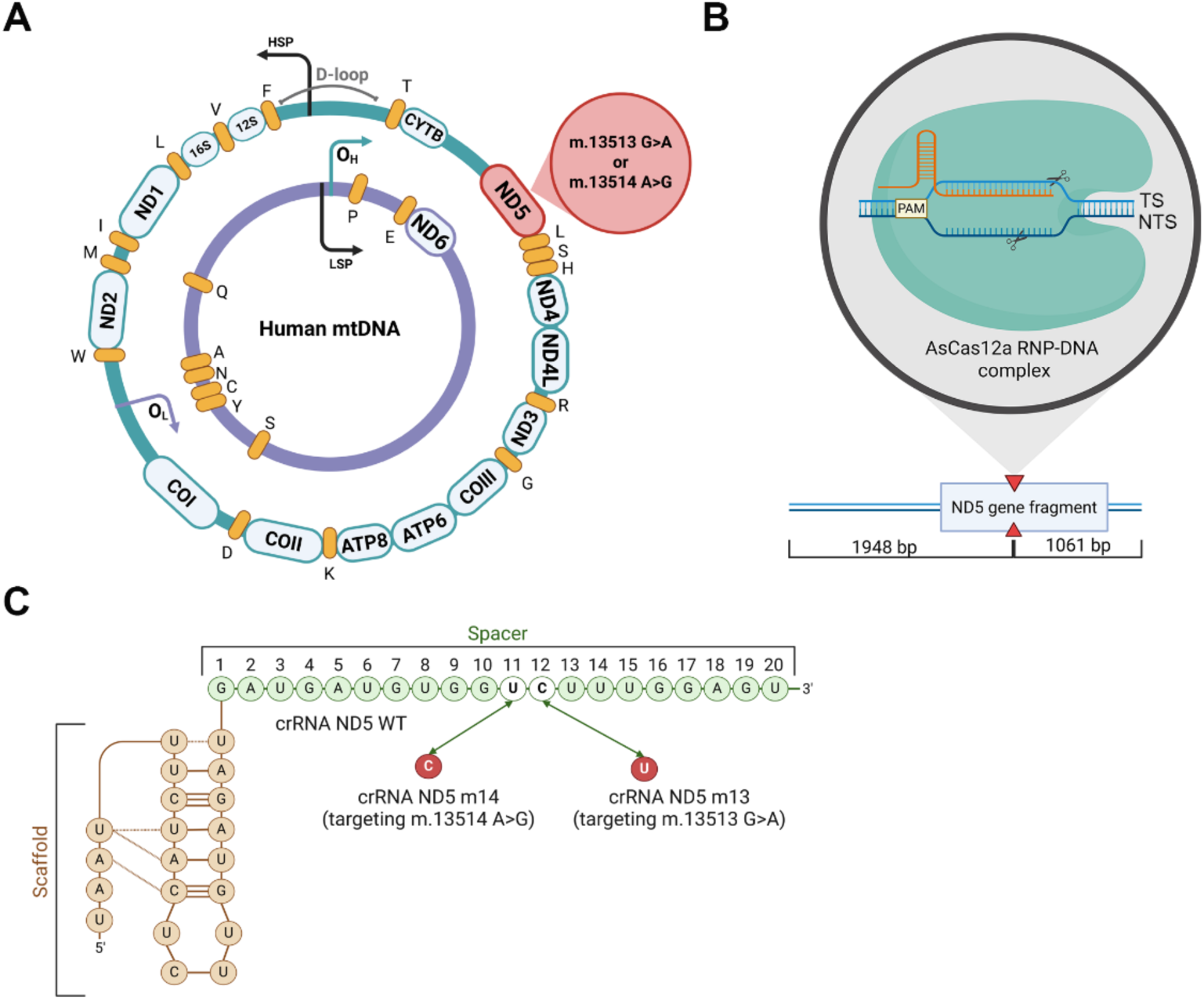
Specific cleavage of mtDNA fragment by AsCas12a effector. **A**, Map of the human mitochondrial genome. Genes are indicated: NADH dehydrogenase subunits (ND1-ND6 and ND4L), cytochrome b (CYTB), cytochrome c oxidase subunits (COI-III), ATP synthase subunits (ATP6, ATP8), rRNA 12S and 16S and tRNAs (denoted by the one-letter code of the corresponding amino acids). HSP and LSP are heavy and light chain promoters; O_H_ and O_L_ are the replication origins of heavy and light chain. D-loop is a non-coding region. Mutations in ND5 gene are highlighted in red. **B**, Schematic representation of Cas12a/crRNA/DNA complex. TS, NTS - target and non-target DNA strands; PAM - protospacer adjacent motif; DNA cleavage sites are indicated. Below, cleavage of the mtDNA fragment used for *in vitro* test is shown. **C**, Predicted 2D structure of crRNA targeting wild-type ND5 gene: nucleotide changes in crRNAs corresponding to mtDNA mutant versions m.13513G>A and m.13514A>G are shown with arrows. Numbers of the nucleotides in the crRNA spacer domain are indicated above.

The natural CRISPR-Cas systems provide adaptive immunity to bacteria and archaea (4). Individual crRNAs, transcribed from CRISPR arrays, program Cas effector nucleases to recognize and cleave nucleic acids containing sequences complementary to crRNA spacers (5). Cas12a, a class 2 Type V-A effector nuclease that recognizes TTTN or TTN protospacer adjacent motif (PAM), possesses a single nuclease domain RuvC, which cleaves two DNA strands in a sequential mechanism, in which initial cleavage of the non-target strand (NTS) is followed by target strand (TS) cleavage (**Fig. 1B**) (6,7). The RNA component of the system, crRNA that programs the target DNA cleavage, consists of a constant 5’ scaffold moiety, which is identical for all crRNAs, and a variable spacer or guide segment, complementary to target DNA (**Fig.1C**) (8,9).

Cas12a effector nuclease has been largely used to edit nuclear genomes in a number of organisms, including human cells (10,11). Eukaryotic nuclear genome editing by CRISPR-Cas is based on the double-stranded breaks (DSB) of DNA in the region of interest, which can be repaired resulting in small insertions/deletions or incorporation of desired template (reviewed in (12)). The possibility of exploiting such convenient and robust system as CRISPR for mitochondrial genome editing is debated due to the low efficiency of guide RNA import into mitochondria (13). Nevertheless, the data from different laboratories show the presence of nuclear-encoded small non-coding RNAs in the mitochondrial matrix (14,15). We suggest that the Type V AsCas12a nuclease is a promising candidate for mitochondrial genome targeting due to its relatively short crRNA (40-nt), which we recently have shown to be importable into human mitochondria (16) and capable of inducing deletion in mtDNA (17). We have also shown that AsCas12a endonuclease can utilize split versions of crRNA, where spacer and scaffold parts added separately, could efficiently cleave target DNA (18). More recently, we reported that RNA import determinants can target human mitochondria with short DNA aptamers permitting mtDNA sequence change by microhomology-mediated mechanism (19).

In contrast to the nucleus, human mitochondria are supposed to lack an efficient double-strand breaks repair systems; it was demonstrated that linear mtDNA molecules after DSB are rapidly eliminated (20), and the total pool of mtDNA molecules then can be restored by replication of wild-type mtDNA molecules (21). Thus, the specificity of mitochondrial genome cleavage, especially discrimination of point mutations, is extremely important to avoid the wild-type mtDNA degradation. The specificity of cleavage by effector nuclease depends on the discrimination capacity of the crRNA spacer region. This is challenging, since several studies reported position-dependent mismatch tolerance within the crRNA by *in vitro* experiments and nuclear DNA editing assays (22–24).

To improve the targeting specificity, enhanced Cas12a nucleases have been designed (25,26). Another approach is to modify crRNAs by engineering its secondary structure (27), introducing additional mismatches or chemical modifications (reviewed in (28)). Recently, chimeric crRNA/DNA molecules were designed, which drastically reduced off-target cleavage (29). Up to now, these studies have been performed on human nuclear DNA sequences.

Here we aimed to study whether AsCas12a/crRNA system is able to discriminate between wild-type and mutant mtDNA molecules harboring pathogenic point mutations in the ND5 gene, coding for the fifth subunit of NADH dehydrogenase, the multi-protein respiratory complex I localized in the inner mitochondrial membrane (**Fig. 1A**) (30). We used as a model mtDNA bearing point mutations in ND5 gene, m.13513G>A and m.13514A>G, associated with multisystem human diseases. Mutation m.13514A>G, detected in patients with MELAS-like and Leigh syndromes, induced amino acid replacement D393G localized in a very conserved region of ND5 transmembrane helix 12 (31). Another frequently reported and well-documented m.13513G>A mutation, associated with a visual loss, deafness and nephropathy (reviewed in (32)) affected also D393 residue in the ND5 gene, but the amino acid replacement, D393N, is different. D393 residue, located in the proton channel, was suggested to be involved in a cascade of conformational changes leading to proton translocation. Therefore, any mutation exchanging the negatively charged D393 to uncharged amino acid residue could disturb the transport of the proton through the ND5 subunit channel of respiratory complex I, which caused a respiratory chain deficiency and clinical manifestations (33).

To improve the specificity of mutant mtDNA cleavage by AsCas12a/crRNA system, we tested different chemically synthetized versions of full and split crRNAs and corresponding chimeric molecules bearing deoxyribonucleotides at various positions of the spacer region. This permitted us to find a crRNA version capable of distinguishing wild-type and mismatched mtDNA sequences under physiological conditions.

Rapid degradation of cleaved mtDNA complicates the detection of targeted double-strand breaks induced by Cas12a system *in vivo*. This issue can be resolved by use of base editors, enzymes which consist of cytosine or adenine deaminases fused to an inactivated version of the AsCas12a nuclease, thus allowing targeted deamination of cytosines or adenines in the region of interest without mtDNA cleavage (34). To check the possible application of the AsCas12a/crRNA to mtDNA in living cells, we designed a mitochondria-targeted cytosine base editor (mtCBE) and demonstrated its low, but significant and specific activity in cultured human cells.

## MATERIAL AND METHODS

### Cloning of ND5 mtDNA fragment for *in vitro* cleavage tests

Genomic DNA isolated from 143B cell line by NucleoSpin Tissue DNA Purification Kit (Macherey-Nagel) was used to amplify a fragment of the ND5 gene of human mtDNA (location 13355-13722 nt) with EcoRI-ND5 and HindII-ND5 primers (**Table S1**) using Phusion High-Fidelity DNA Polymerase (Thermo Scientific). The amplified fragment was cloned into the pUC19 vector. The resulting plasmid, named pUC19_ND5_WT, was subsequently mutated to introduce either the 13513G>A or 13514A>G mutation *via* site-directed mutagenesis using Q5 Site-Directed Mutagenesis Kit (NEB) with primers ND5 mut m13 or ND5 mut m14 and ND5 mut R (**Table S1**). For *in vitro* cleavage tests, the plasmids were linearized with the ScaI (Thermo Scientific).

### *In vitro* DNA cleavage assay

All crRNAs used in this study are listed in **Supplementary Table S1**. To design crRNA targeting the pathogenic mutations 13513G>A and 13514A>G, ND5 gene of mtDNA (GenBank NC_012920.1, gi:251831106) was analyzed for the sequences representing the canonical AsCas12a "TTTV" PAM (35) near the region of interest. The chosen 20 bp-long sequences of spacers in the 5’ to 3’ direction were supplemented at the 5’ end by the AsCas12a scaffold sequence (36). These crRNAs were then analyzed to predict the probability of introducing unintended changes (off-target effects) by the Cas-OFFinder algorithm (37). The secondary structures of AsCas12a crRNAs were predicted using the RNAfold web server (38) and the Mfold web server (39).

The chemical synthesis of crRNA molecules was performed as described previously (40). Briefly, crRNAs were synthesized on an automatic ASM-800 RNA/DNA synthesizer (Biosset) using phosphoramidites purchased from Glen Research. After standard deprotection, crRNAs were purified *via* preparative electrophoresis in 12% polyacrylamide/8 M urea gel and characterized by MALDI-TOF mass spectrometry (Autoflex III, Bruker Daltonics). The recombinant AsCas12a nuclease was purchased at abm (K188).

*In vitro* cleavage assays were carried out in 20 mM HEPES (pH 7.5), 100 mM KCl, 1 mM DTT, 5% glycerol, MgCl_2_ (1 mM, 2 mM, or 5 mM) and 0.1% Triton-X100 (if required, according to experimental setup). The standard reaction contained 2.9 nM DNA templates, 133.3 nM AsCas12a and 73.4 nM of 40-nt crRNA or 146.8 nM of split crRNA. Reactions were incubated for 30 min at 37°C and stopped by the addition of 20 *μ*g of Proteinase K (Promega) at 37°C for 10 min. Analysis of cleavage products was performed by electrophoresis in 1% agarose gel in 1× TAE buffer. EtBr-stained gels were visualized by the Hero Lab Transilluminator UVT- 28M (Herolab GmbH) and the DNA fragments were quantified with ImageJ Software. Cleavage efficiency was calculated as % of cleaved DNA fraction; for statistical analysis of cleavage efficiency, two-tailed unpaired t-test was used.

### UV-Melting Experiments

Thermal denaturation assays of the crRNA/DNA duplexes were carried out in an equimolar mixture containing 2.5 µM of each strand in a buffer containing 50 mM Tris-HCl, pH 8.0, 60 mM NaCl, and 1 mM MgCl_2_, as described in (41). DNA oligonucleotide sequences are shown in **Table S1**. Melting curves were recorded on a Cary 300 Bio UV-VIS spectrophotometer (Varian) equipped with a Peltier thermostabilized multicell holder at wavelengths 260, 270, and 330 nm in the 5−90°C range at a temperature change rate of 0.5°C/min. Absorbance at 330 nm was subtracted from the values at other wavelengths at each temperature. Tm was calculated as a maximum of the first derivative of optical absorbance in terms of temperature. The data (**Table S2**) represent an average from three experiments on denaturation and renaturation. The experimental error Tm is ±0.2°C.

### Mammalian cell culture

HEK293 cell line and its derivatives were maintained at 37°C with 5% CO₂ in essential modified Eagle’s medium (EMEM, Sigma Aldrich), which contained 1 g/L D-glucose, supplemented with 1.5 g/L sodium bicarbonate (Sigma Aldrich), 0.11 g/L sodium pyruvate (Sigma Aldrich), 10% fetal bovine serum (Gibco, Fisher Scientific), 100 mg/L penicillin-streptomycin (Sigma Aldrich), and 2.5 mg/L amphotericin B (Sigma Aldrich).

Before each experiment, cell cultures were tested for mycoplasma contamination. The presence of Mycoplasma was tested using PCR with DreamTaq Green PCR Master Mix (Thermo Scientific), as described by (42) (**Table S1**). Total DNA was isolated using the NucleoSpin® Tissue Mini Kit (Macherey Nagel), following the manufacturer’s protocol, and used as the PCR template.

### Assembly of mtCBE plasmid

To assemble mitoCBE-AsCas12a base editor (mtCBE) we used Su9 MTS and 3xFLAG epitope tag from our lab collection (17) and cytidine base editor (CBE, APOBEC3) fused to the N-terminus of enAsCas12a (23), inactivated by a D908A mutation (denAsCas12a)(Addgene Plasmid #171698). Uracil-DNA glycosylase inhibitor (UGI) was attached to the C-terminal end of the base editor. Resulting construct was cloned into pET30a (+) (Novagen) for protein purification and into pcDNA5/FRT (Thermo Scientific) for transfection of cultured cells.

### Expression and purification of recombinant mtCBE

The mitoCBE-AsCas12a base editor (mtCBE, **Fig. S3 A**) cloned to pET30a (+) vector was used for transformation of *E. coli* expression strain Rosetta (DE3) (Novagen). Then plasmid-containing bacterial colony was picked and cultivated at 37°C in 100 ml of LB medium, supplemented with 100 *μ*g/mL of ampicillin, until the optical density at 600 nm (OD600) reached 0.8. Subsequently, the temperature was lowered to 18°C. Nuclease expression was induced by adding isopropyl-*β*-D-1-thiogalactopyranoside (IPTG) to a final concentration of 500 *μ*M. The culture was then maintained at 18°C for 12-16 hours. The cell pellet was collected by centrifugation at 8,000 g for 10 minutes at 4°C and then resuspended in 1x PBS (137 mM NaCl, 2.7 mM KCl, 10 mM Na2HPO4, and 1.8 mM KH2PO4), supplemented with 1x Protease Inhibitor Cocktail (Roche). Subsequently, the cells were lysed by several rounds of sonication for 10 seconds, using a Vibra-Cell™ VCX 750 Sonicator (2 mm probe) at a frequency ranging from 20 to 50 kHz with intervals of 1 minute, on ice. The resulting lysate was centrifuged at 14,000 g for 20 minutes at 4°C. The supernatant was then incubated with Ni-NTA agarose (Qiagen) according to manufacturer’s recommendations. The protein of interest was eluted with 1xPBS and 250mM imidazole, concentrated, snap-frozen and stored at -80°C.

### *In vitro* deamination tests

*In vitro* deamination assays were carried out in Reaction buffer (20 mM HEPES (pH 7.5), 100 mM KCl, 1 mM DTT, 5% glycerol and 1 - 5mM MgCl_2_) (43,44). The DNA templates, representing the fragments of human ND4 and ND5 genes, were obtained by PCR amplification with Phusion High Fidelity DNA Polymerase, purified by NucleoSpin Gel and PCR Clean-up Mini Kit (Macherey-Nagel) and the DNA concentration was measured by NanoDrop 1000 Spectrophotometer V3.5 (Thermo Scientific). The standard reaction contained 300 ng of DNA template, 4 *µ*g of mtCBE protein and 400 ng of 40-nt crRNA or 200 ng of spacer and 200 ng of scaffold for split crRNA. Reactions were incubated for 24 hours at 37°C and stopped by the addition of 20 *μ*g of Proteinase K (Promega) and incubation at 37°C for 10 min. The reactions were purified with NucleoSpin Gel and PCR Clean-up Mini Kit (Macherey-Nagel) and sent for Sanger sequencing. The results of *in vitro* deamination were analyzed by R-based software EditR (45) and are shown in **Supplementary Table S3**.

### Immunofluorescent staining followed by confocal laser scanning microscopy

All antibodies used in this study are listed in **Supplementary Table S4**.

Immunofluorescent staining was performed following the method described in (46), with modifications. Briefly, cells were seeded at a density of 0.15–0.20 cm² in 8-well Nunc Lab-Tek slides (Thermo Fisher Scientific) pre-treated with 0.1% gelatin for 1 hour at 37°C. After 24 hours, the cell culture medium was replaced with 200 µL of freshly preheated EMEM at 37°C. The transfection mix was prepared by combining 100 ng of plasmid DNA (**Fig. S3 C**) with 0.5 µL of Lipofectamine™ 2000 Transfection Reagent (Invitrogen) in 100 *µ*L of Opti-MEM™ (Gibco, Fisher Scientific), following the manufacturer’s protocol, and added to the plate. Cells were allowed to propagate for an additional 24 hours.

For immunofluorescence staining, cells were fixed in a preheated fixation buffer (4% paraformaldehyde in PBS, adjusted to 3% using EMEM) for 12 minutes at 37°C. After fixation, the cells were washed five times with 1× PBS. Next, cells were permeabilized using a permeabilization buffer (0.5% Triton X-100 in 1× PBS) and incubated for 15 minutes at room temperature (RT). After another round of washing, the cells were blocked with a blocking buffer (5% BSA in 1× PBS). Following the incubation, the blocking solution was removed, and primary antibodies (**Table S4**) diluted in the blocking buffer were added. The cells were incubated with the primary antibodies for 3 hours at RT. After washing, secondary antibodies conjugated with a fluorescent marker were diluted in the blocking buffer and added to the cells for 1 hour at RT. From this step onward, all incubations were performed in the dark.

After the final wash, nuclei were stained by incubating the cells with 1 µg/mL of DAPI (Invitrogen) in 1× PBS for 10 minutes at RT, followed by washing.

The samples were imaged using an LSM700 microscope (Carl Zeiss) with a 63×/1.4 oil objective in 1× PBS. Images were visualized and analyzed using ImageJ 1.52a software (47).

### *Ex vivo* deamination test – cell transfection with RNA

HEK293 cells (2 cm^2^) were transfected in suspension with 500 ng mtCBE-expressing plasmid and 300 ng of crRNA using Lipofectamine 2000 transfection reagent (Invitrogen) in Opti-MEM (Gibco, Fisher Scientific), twice with three days interval. Two days after second transfection, the cells were detached, and total DNA was isolated using the QIAamp DNA Mini Kit (Qiagen) for further NGS analysis.

### NGS libraries preparation

Fragments of mtDNA obtained by PCR using either Phusion High-Fidelity DNA Polymerase (Thermo Scientific) or DreamTaq Green PCR Master Mix (Thermo Scientific) were purified with the NucleoSpin Gel and PCR Clean-up Mini kit (Macherey-Nagel) according to the manufacturer’s protocol. DNA concentrations were measured using a NanoDrop 1000 Spectrophotometer (Thermo Scientific).

For library preparation, 100 ng of isolated DNA was used following the NEBNext Ultra II FS DNA Library Prep Kit for Illumina (NEB). Adapters and indexes were provided by the NEBNext Multiplex Oligos for Illumina (Dual Index Primers Set 1) (NEB). During the library preparation process, DNA purification was carried out using the NucleoMag NGS Clean-up and Size Select kit (Macherey-Nagel) and the NucleoMag SEP Mini separation plate (Macherey-Nagel).

Quality control of the libraries, including Bioanalyzer and Qubit measurements, along with sequencing, was conducted at the Plateforme d’analyse de l’expression génique (IBMP, Strasbourg). Sequencing was performed on a MiSeq (Illumina) system, generating 150-bp paired-end reads with a target coverage of approximately 1,000,000 reads per sample.

The obtained reads were analyzed using the Galaxy server (version 23.1.rc1) (48). Nucleotides with a quality score below 20 were filtered out using Trimmomatic (version 0.38) (49). The filtered reads were mapped to the reference mtDNA genome (GenBank NC_012920 gi:251831106) using BWA-MEM2 (version 2.2.1) (50). NCBI BLAST software was used to predict possible NUMTs and, if presented, to exclude them from the analysis.

The resulting alignments were visualized with IGV (version 2.16.1) (51). Further processing of the sequencing data was done using BAMtools (version 2.5.1) (52) and SAMtools (version 1.13) (53). For statistical analysis of nucleotide substitutions quantification, two-tailed unpaired t-test was used.

## RESULTS

### Modified crRNAs provide specific cleavage of mtDNA by AsCas12a

To assay the possibility of mutant mtDNA specific cleavage by Cas12a/crRNA, we used as a model mtDNA bearing two point mutations in ND5 gene, m.13513G>A and m.13514A>G, further referred to as m13 and m14 correspondingly(31). We designed crRNA targeting the region of interest (**Fig. 1C**); notably, due to position of the nearest PAM sequence (TTTN) in mtDNA, the mutated nucleotides are located in PAM-distal positions 11 and 12. This so-called “mid-region” of the crRNA spacer was previously characterized by incomplete mismatches discrimination (28,54). Therefore, we addressed the question whether specific cleavage of mutant mtDNA molecules can be performed by the Cas12a/crRNA system designed. For this, human mtDNA wild-type or mutant sequences were cloned and used as a target for *in vitro* cleavage tests in presence of recombinant AsCas12a nuclease and chemically synthetized crRNAs (18). First, we performed cleavage under standard conditions optimized previously for Cas9 and Cas12a nucleases activity measurement (18,55,56), using combinations of three crRNAs (WT, m13 and m14) and three DNA target sequences (**Fig. 2A**). The data showed that, indeed, the system demonstrated a low level of discrimination between wild-type mtDNA sequence (DNA WT) and mutant versions (DNA m13 and m14), especially for crRNAs designed to target mutant DNA (shown in green and orange, **Fig. 2A**), which are able to induce the cut of WT mtDNA almost at the same level as for cognate mutant DNA (90-95%).

**Figure 2:**
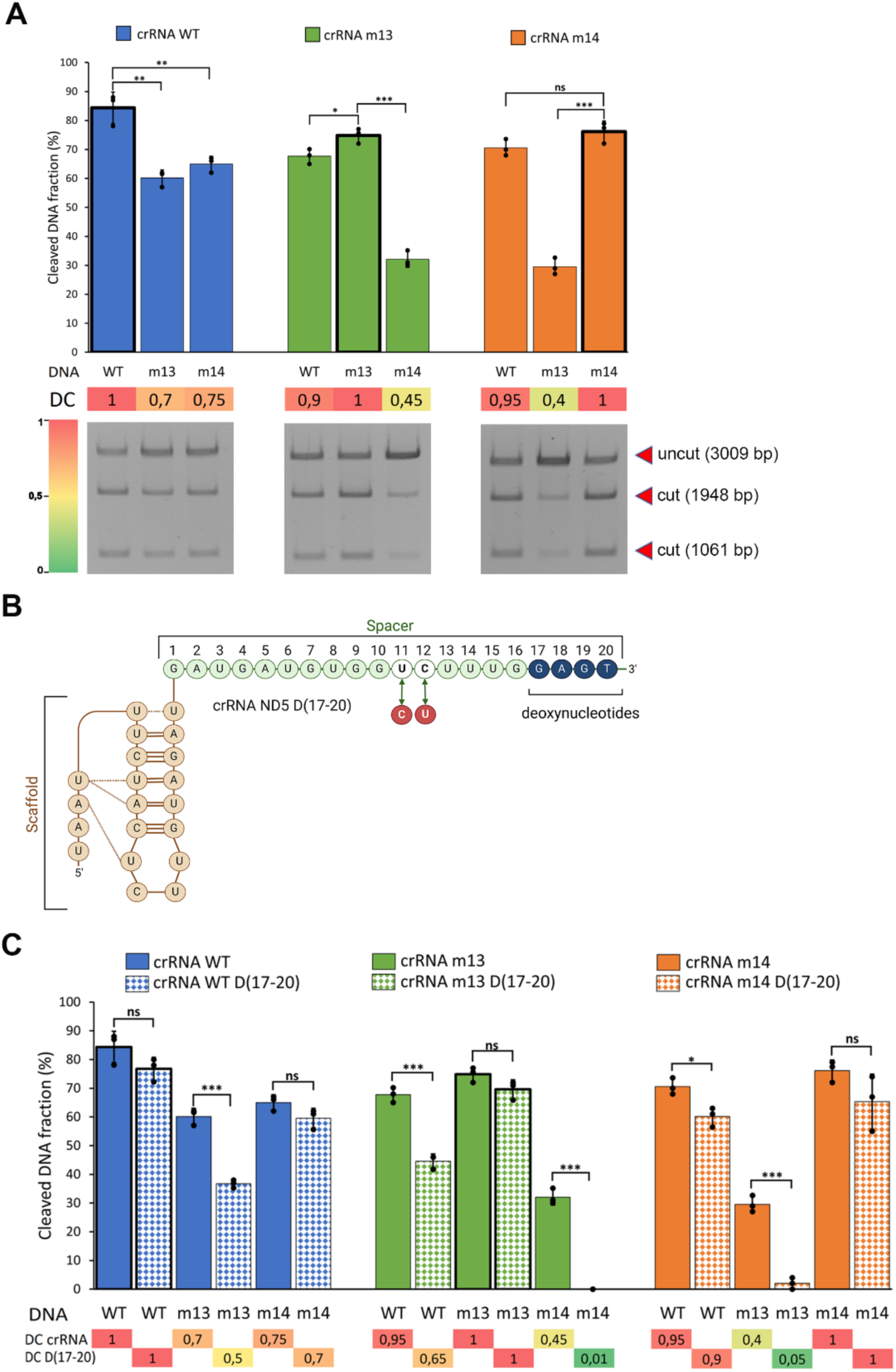
Improvement of cleavage specificity by chimeric crRNA D(17-20). **A**, *In vitro* cleavage test of target DNA (WT, m13, m14 as indicated below the panels) by AsCas12a effector programmed with crRNAs indicated above the panels, at 5 mM MgCl_2_; n=3, representative agarose gel is shown on the lower panel, DNA fragments size indicated at the right. DC, Discrimination Coefficient calculated as a ratio between cleavage efficiency (% of cleaved DNA fraction) for mismatched and cognate DNA/crRNA pairs. Number of replicates = 3. **B**, Sequence and structure of crRNA D(17-20). **C**, Comparison of the target DNA cleavage by AsCas12a effector programmed with non-modified crRNAs and crRNAs D(17-20), shown by solid and dashed coloring correspondingly, at 5 mM MgCl_2_. Discrimination coefficients are shown and colored according to the heat map. Discrimination coefficient was estimated for each non-modified crRNA (upper line) and crRNA D(17-20) independently, as indicated. Number of replicates = 3.

To improve the cleavage specificity, we introduced four deoxynucleotides in the distal region of the crRNA part (**Fig. 2B**), referred to as D(17–20) (25,29). The cleavage test demonstrated increased specificity for versions bearing two mismatches (as for crRNAm13/DNAm14), but only a slight improvement of the discrimination for point mutations (**Fig. 2**, **Suppl. Fig. 1A**).

Then, the technology of split crRNA, which we developed previously (18), was applied. We had shown that Cas12a effector proteins can be programmed by crRNA split into two parts, corresponding to the scaffold, derived from the structured direct repeat, and the spacer region, complementary to the DNA target (**Fig. 3**). This strategy facilitates the chemical synthesis of modified crRNA, because different 20-nucleotide spacer domains can be associated with the same scaffold RNA. Therefore, we have tested a set of spacer sequences targeting WT, m13 and m14 DNA, bearing deoxynucleotides in various positions (**Fig. 3**, **Suppl. Fig. 1B**, **Table S1**). Cleavage assays demonstrated that the version carrying eight deoxynucleotides in the distal region of the crRNA spacer (D13-20) could not program any cleavage of the DNA template (**Suppl. Fig. 1B**). Split crRNA with four deoxynucleotides D(17–20) was characterized by significantly improved discrimination capacities compared to full D(17–20) crRNA versions (**Fig 3A**), thus showing, for the first time, the potential to use split crRNAs for more specific DNA cleavage and editing. Surprisingly, split crRNA bearing four deoxynucleotides in positions 13-16 of the spacer region demonstrated best ability to distinguish between wild-type and mutant mtDNA versions, since the cleavage of non-cognate DNA targets was very low or not detected (**Fig. 3B**, **Suppl. Fig. 1C**). At the same time, the cleavage efficiency of cognate targets m13 and m14 by split crRNAs D(13–16) was also slightly reduced compared to split crRNAs D(17–20).

**Figure 3:**
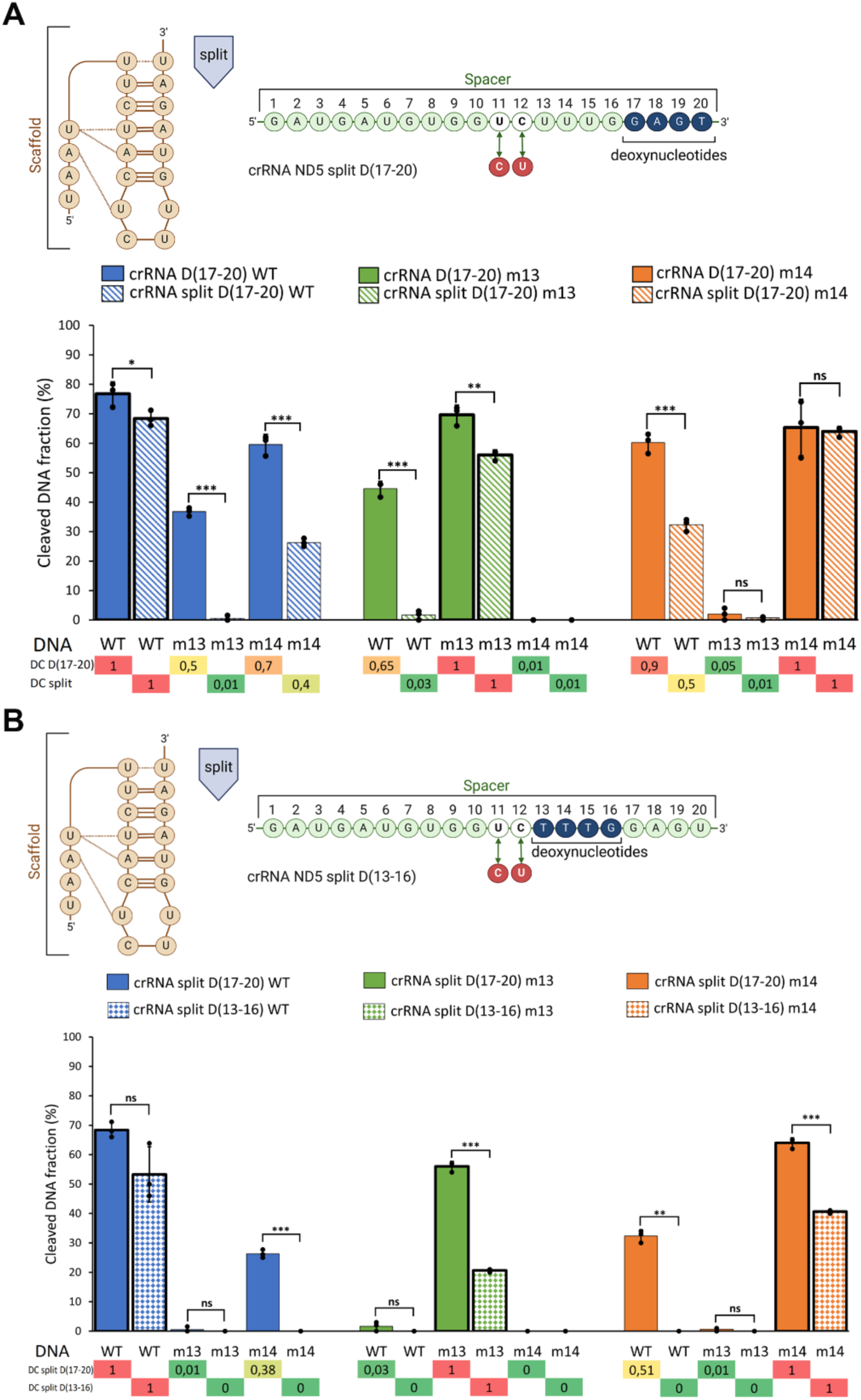
Effect of split crRNA modifications on the cleavage specificity at 5 mM MgCl_2_. **A**, Structure of split crRNA D(17-20) (upper panel) and comparison of the DNA cleavage by AsCas12a effector programmed with crRNAs D(17-20) and split crRNAs D(17-20) versions, shown by solid and dashed coloring. Number of replicates = 3. **B**, Structure of split crRNA D(13-16) and DNA cleavage induced by split crRNA versions D(17-20) and D(13-16), as indicated. Number of replicates = 3.

Therefore, we identified crRNA modifications improving the ability of the AsCas12a/crRNA system to cut only the target DNA without mismatches, at least under conditions used.

### Physiological conditions improve the specificity of cleavage

It is questionable if the conditions described above match the situation *in vivo*, notably inside human mitochondria. From the beginning of the CRISPR-Cas systems studies, *in vitro* and in cell lysate cleavage tests have been performed in buffers with rather high (5 – 10 mM) MgCl_2_ concentrations, which assured maximal nuclease activity (7,54–56). Noteworthy, intercellular concentrations of free Mg^2+^ ions in bacteria (57), eukaryotic cells and mitochondria (58) have been estimated as 0.5 – 1.2 mM. Moreover, the *in vitro* reactions mixtures often contained detergent Triton X-100 (8,18,55), thus creating conditions quite different from the *in vivo* ones. Taking into account this discrepancy, we compared AsCas12a/crRNA cleavage efficiency in buffers containing different Mg^2+^ concentrations with and without detergent, using a cognate wild-type crRNA-DNA WT combination as well as non-cognate, namely crRNA WT with mutant mtDNA templates m13 and m14 (**Fig 4**). For all the DNA templates, we demonstrated a significant increase of nuclease activity upon increase of Mg^2+^ concentrations from 1 to 5 mM, as it was reported previously (59). Surprisingly, presence of 0.1% Triton X-100 also increased the cleavage level, however the difference was mostly non-significant.

**Figure 4:**
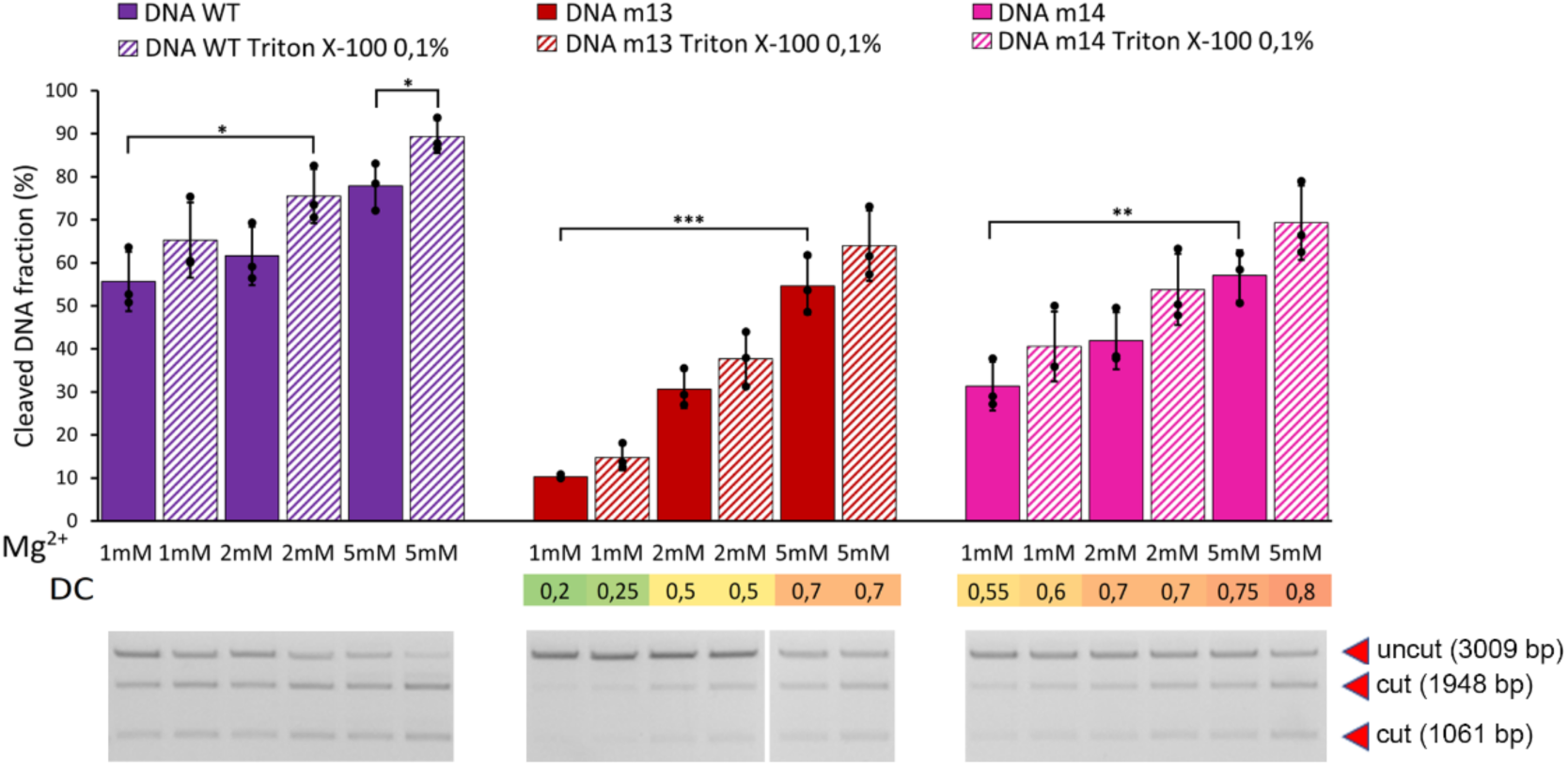
AsCas12a/crRNA cleavage efficiency in buffers containing different Mg^2+^ concentrations. Cleavage of different target DNA (indicated above the panels) by AsCas12a effector programmed with crRNA WT at Mg^2+^ concentrations from 1 to 5 mM, as indicated below the panels, in presence (dashed coloring) or not of detergent Triton X-100. Number of replicates = 3.

Noteworthy, discrimination coefficients (ratio between cleavage of cognate and non-cognate DNA templates) were not dependent on the presence of 0.1% Triton X-100, but dramatically changed at lower Mg^2+^ concentration, decreasing from 0.7-0.8 at 5 mM Mg^2+^ to 0.2-0.5 at 1 mM Mg^2+^ (**Fig. 4**). Interestingly, at low magnesium, we obtained higher tolerance to m.13514A>G mutation (m14, corresponding to mismatch U:G in position 11 of crRNA spacer) compared to m.13513G>A (mismatch C:A in position 12), the difference which perfectly corresponds to the melting temperatures (Tm) of target DNA/ crRNAs duplexes measured by thermal denaturation assay (**Table S2**).

Therefore, we suggested that under physiological conditions, DNA cleavage by AsCas12a/crRNA can be less efficient but more specific. To validate our data and to further optimize the mismatch discrimination, we compared the specificity of DNA cleavage by AsCas12a programmed with full, split and DNA-modified crRNAs at 1 mM Mg^2+^. For full gRNAs in low magnesium, D(17–20) modification greatly improved the cleavage specificity, reducing the cut of mismatched gRNA/DNA pairs from 0.5-0.9 (**Fig. 3A**) to 0.1-0.3 (**Fig 5, Suppl. Fig. 2A**). For split crRNA versions, surprisingly, variant D(13–16), characterized by the best discrimination abilities in 5mM Mg^2+^ (**Fig. 3B**), could not program any DNA cleavage in 1 mM Mg^2+^ (**Suppl. Fig. 2C**). In contrast, crRNA split variant D(17–20) demonstrated best ability to distinguish between wild-type and mutant mtDNA versions (**Fig 5**, **Suppl. Fig. 2B**).

**Figure 5:**
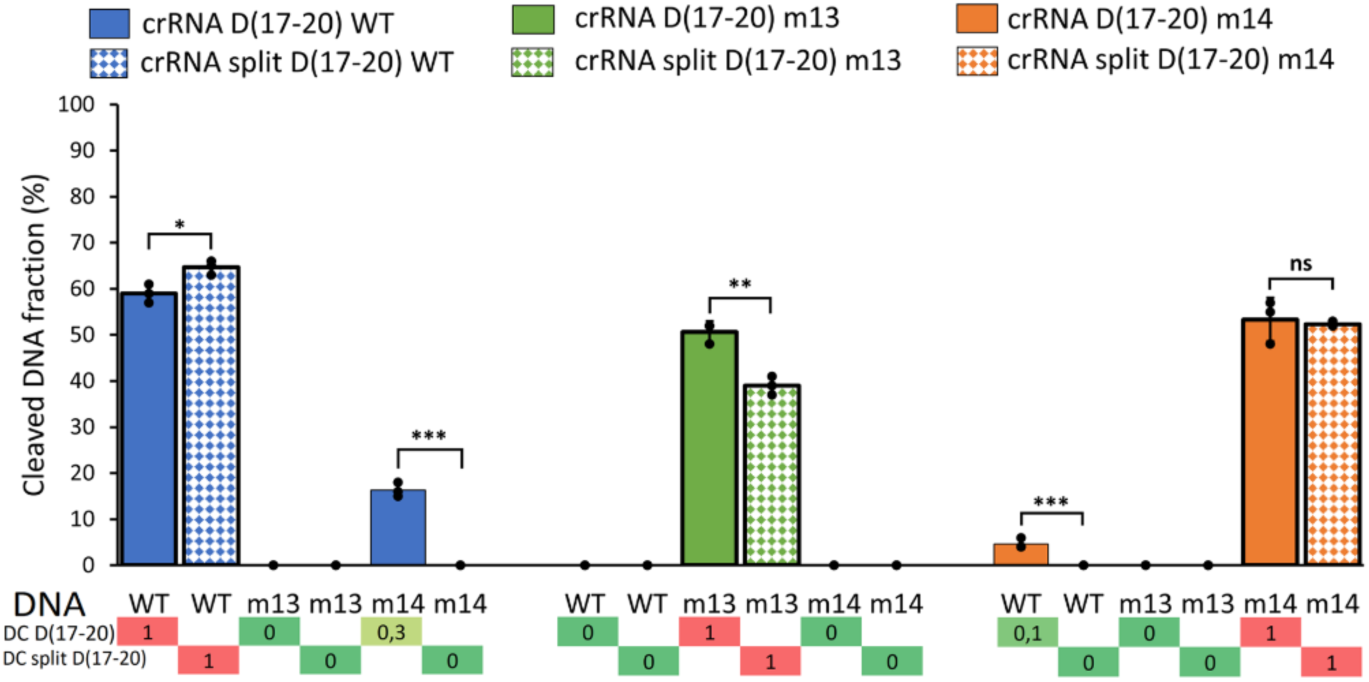
Effect of crRNAs bearing deoxynucleotides on the cleavage specificity at 1 mM MgCl_2_. Target DNA cleavage by AsCas12a effector programmed with crRNAs D(17-20) and split crRNAs D(17-20) version (shown by solid and dashed coloring) at 1 mM MgCl_2_. Number of replicates = 3.

In conclusion, our data show that the mismatches in positions 11 and 12 of the Cas12a crRNA spacer region can be tolerated under high Mg^2+^ concentration and in presence of detergent. At low Mg^2+^, corresponding to its concentration in cells and mitochondria, the discrimination abilities of AsCas12a/crRNA system were significantly improved. Split crRNAs, especially their modified versions bearing four deoxynucleotides in the distal region of the spacer part, D(17–20), provided best discrimination properties.

These findings will be helpful for further *ex vivo* experiments, aiming to obtain specific cleavage of mutant mtDNA in *trans*-mitochondrial cybrid cell lines and in patients’ primary fibroblasts bearing pathogenic mutations m.13513G>A and m.13514A>G, which can shift the load of the mtDNA mutation and thus improve the mitochondrial functions.

### Setting up mitochondrial Cas12 base editing system in living cells

Due to unique features of mitochondrial DNA maintenance, cleaved mtDNA is rapidly degraded by components of the mitochondrial genome replication machinery (20). This complicates the detection of targeted double-strand breaks and the assessment of the mitoCas12a efficiency in living cells (*ex vivo*). This problem can be solved by using base editors ((26), reviewed in (60)), which induce single-nucleotide substitutions that can be further detected by DNA sequencing.

For this *ex vivo* application, we used cytosine deaminase CBE (APOBEC3) allowing targeted deamination of cytosines (61) fused to an inactivated version of the AsCas12a nuclease (dCas12a). The dCas12a protein guided by crRNA binds to and unwinds the target DNA sequence, where CBE catalyzes deamination of Cs to Us in the region of NTS (non-target strand) called editing window comprising approximately 7 nucleotides.

To assure the import of the base editor into mitochondria, N-terminal peptide MTS (mitochondrial targeting signal) Su9 was added to the construct (**Fig. S3A**) (17). Resulting protein, referred to as mtCBE, can be expressed in cultured human cells and co-localized with mitochondria (**Fig. S3B**, **C**).

First, we performed *in vitro* deamination test with recombinant mtCBE and crRNA m14 aiming to correct the ND5 m.13514A>G mutation back to the wild-type by deamination C on the opposite strand, thus inducing replacement of G13514 to A. Since the base editors target a window of about 7 nucleotides, unwanted deamination (bystander editing) may occur on several C residues (62).Indeed, we detected the simultaneous deamination of two cytosines within the editing window (**Fig. S3D**). Thus, we obtained a partial replacement G13514 to A, but also introduced the m.13513G>A mutation. Modified D(17–20) crRNA m14 induced deamination compared to the unmodified crRNA, indicating that modifications didn’t impact the base editing (**Fig.S3D**, bottom panel). We also checked the level of *in vitro* editing at increased concentrations of MgCl_2_ (**Fig. S4**). In contrast to the nuclease activity of crRNA/AsCas12a system, which was increased at higher magnesium, we observed a reduced deamination activity of AsCas12a-fused base editor mtCBE at 5 mM MgCl_2_ compared to 1 mM MgCl_2_ (**Fig. S4, Table S3**).

Next, we tested the ability of crRNA WT to introduce the m.13513G>A mutation into wild-type ND5 sequence bearing only one G in the editing window. We compared activity of different crRNAs: unmodified crRNA WT, crRNA WT D(17-20) and split WT crRNA versions (**Fig. 6A**). Sanger sequencing data confirmed that the m.13513G>A mutation was successfully introduced into the wild-type mtDNA sequence *in vitro* by all crRNAs versions tested, with mutation rates of 60-70%. This led us to consider applying mtCBE *ex vivo*, targeting the wild-type ND5 in HEK293 cells with the goal of inducing the pathogenic m.13513G>A mutation. This approach would not only allow us to evaluate the deamination efficiency of the AsCas12a-fused base editor mtCBE system in a mammalian cell line, but also eventually create a human culture cell model harboring heteroplasmic m.13513G>A mutation.

**Figure 6:**
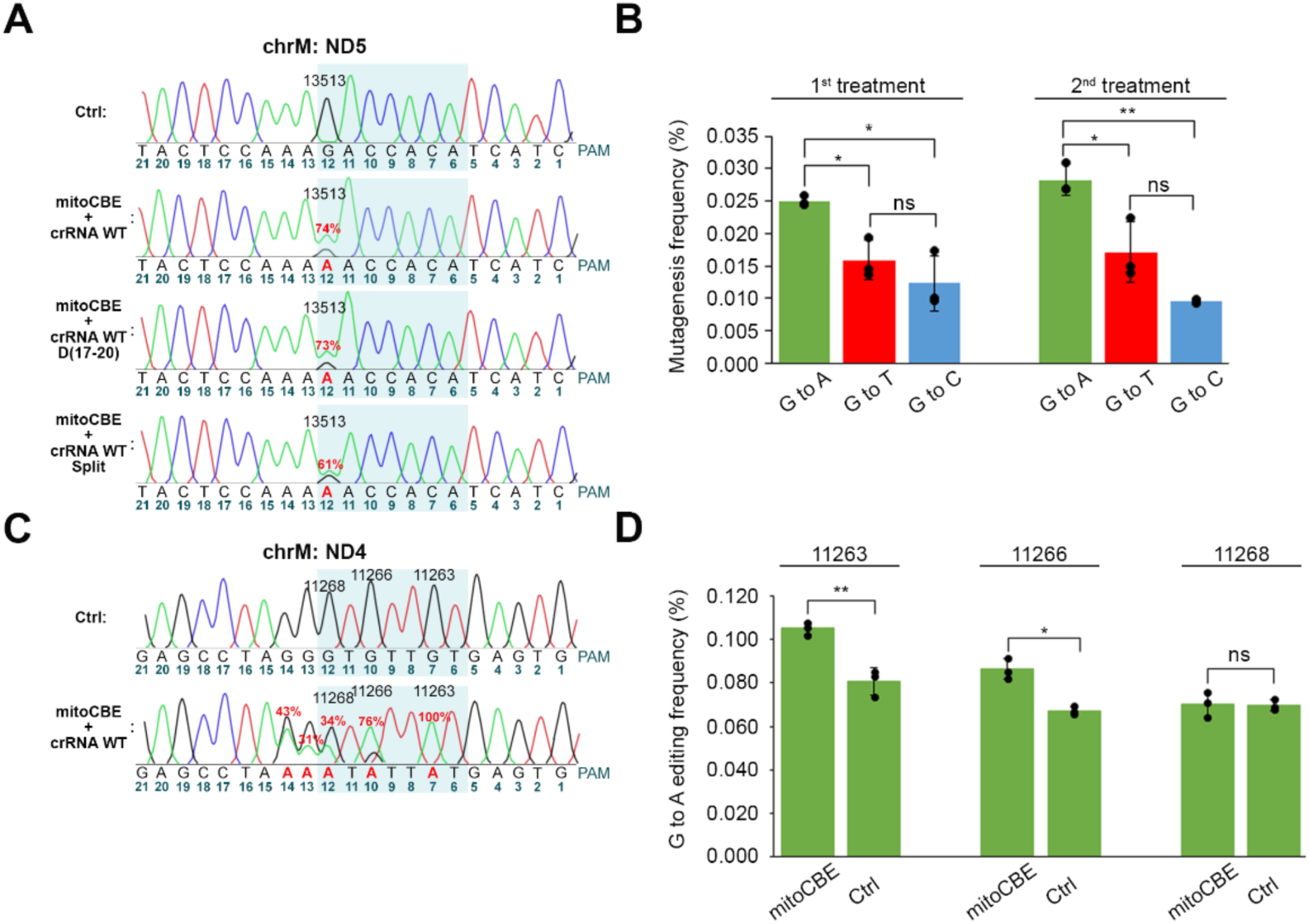
Application of mitoCBE-AsCas12a to HEK293 mtDNA *in vitro* and *ex vivo*. **A**, **C**, Sanger sequencing analysis of a PCR-amplified mtDNA fragment treated *in vitro* with recombinant mitoCBE-AsCas12a in complex with various crRNAs WT indicated at the left, in buffer containing 1mM Mg^2+^. The editing window is highlighted in blue, nucleotide numbers from PAM are below. **A**, The percentage of m.13513G>A mutation introduced by the base editor is indicated in red; the data of one from 2 or 3 experiments (**Table S3**) is shown. **B**, Quantification of nucleotide substitutions (mutagenesis frequency) at position 13513 in the ND5 gene of mtDNA. HEK293 cells were transfected with plasmid expressing mtCBE and crRNA WT, the data after one and two consecutive transfections are shown. Standard deviation is presented for three independent biological replicates. **C**, as in **A**, mitoCBE-AsCas12a was charged with crRNAs targeting the ND4 mtDNA gene to introduce G>A mutations at positions 11263, 11266, and 11268. **D**, Quantification of targeted G to A editing for three nucleotides in the editing window in the ND4 gene of mtDNA in HEK293 cells expressed mitoCBE-AsCas12a and transfected with crRNA targeting this site. MtDNA nucleotide numbers are shown above the graphs. Standard deviation is presented for three independent biological replicates.

For this, HEK293 cells expressing mtCBE were transfected with ND5 WT crRNA, twice with three days interval. Cells after the first and second transfections were used for total DNA isolation, then mtDNA ND5 fragment was amplified and analysed by MiSeq deep sequencing. Analysis of the data demonstrated a low level of G>A editing in transfected cells, however, adenine appeared at position 13513 significantly more often than thymine or cytosine (**Fig. 6B**). This indicates that the level of targeted deamination of cytosine with mtCBE system, while low, is still significantly higher than the rate of random mutations.

The low rate of 13513G>A mutation may be due to the fact that targeted deamination of cytosine occurs at position 12 from the PAM, at the very end of the editing window (**Figs. 1B and 6A**). To verify this hypothesis, we applied mtCBE in complex with a crRNA targeting another region of human mtDNA, namely ND4 gene (14,16). For this crRNA, the editing window covers nucleotides 11262-11268 of mtDNA harboring three Gs at positions 7, 10 and 12 from PAM (**Fig. 6C**). *In vitro* editing test resulted in complete (100%) G>A substitution at position 7 and slightly less for G10 and for G12; moreover, mtCBE additionally deaminated two cytosines outside the expected editing window, at positions 13 and 14 (**Fig. 6C**).

Surprisingly, *ex vivo* editing in HEK293 cells transfected with the same crRNA ND4 resulted in a low but significant C-to-T editing at positions 7 and 10 (compared to the control non-transfected cells) (**Fig. 6D**), and the absence of nucleotide substitution in position 12 (11268). This corroborated our hypothesis regarding the low level of editing at the end of the window.

To summarize this part of the study, we designed a mitochondrial cytosine base editor fused to AsCas12a effector protein and demonstrated that modified crRNAs, namely D(17-20) and split versions can successfully target mtCBE to the template DNA. The system was successfully applied to edit the ND4 and ND5 genes of mtDNA in HEK293 cells. To our knowledge, this is the first reported case of Cas12a-fused base editing of mitochondrial DNA in mammalian cells.

## DISCUSSION

Taken together, our data reveal some important points: 1) split chimeric crRNA substantially improve AsCas12a mismatch intolerance; 2) magnesium concentration has a direct effect on the ability of crRNAs to discriminate single-nucleotide mtDNA variants; and 3) mitochondrially targeted AsCas12a fused to a cytosine base editor enables detectable sequence-specific editing of mtDNA in human cells.

### Modified crRNAs improve the mismatch intolerance of AsCas12a

For all known Cas effectors, the stability of the duplex between crRNA and target DNA determines the efficiency of on-target cleavage. However, some mismatches can be tolerated, which results in off-target cleavage and thus decreases the genome editing accuracy (63,64). CRISPR–Cas12a system was shown to be more sensitive to mismatches between the target DNA and the crRNA than CRISPR–Cas9, the tolerance depending on the position of the mismatch in the crRNA spacer region. In case of Cas12a, when a mismatch was introduced into the seed sequence (5-10 base from the PAM), the cleavage activity was significantly reduced (64). In contrast, off-target cleavage with a mismatch in the PAM-distal region or in the middle of the target sequence was frequently observed (29,63). To enhance the target specificity, chimeric crRNA/DNA molecules were designed, bearing deoxynucleotides at the 3’ end of spacer region, which drastically reduced off-target cleavage (29). It was suggested that deoxynucleotides should decrease the stability of the mismatched DNA/crRNA duplex, but not of the perfect duplex, thus only slightly reducing the target DNA cleavage and strongly affecting off-target cleavage (25).

We applied a similar approach to increase the specificity of Cas12a system toward point mutations in the mitochondrial ND5 gene. Our data demonstrated an improvement in the ability to discriminate mutations by use of chimeric crRNA molecules, which was, as expected, less pronounced for m.13514A>G mutation (corresponding to a weak mismatch U:G in position 11 of crRNA spacer) compared to m.13513G>A (mismatch C:A in position 12) (**Fig. 2**), indicating on the impact of the nucleotide sequence which determines the stability of DNA/crRNA duplex. More spectacular effect was obtained by use of split crRNA, where two crRNA domains, scaffold and spacer, were synthetized and added separately (**Fig. 3**).

To get a deeper understanding of the mechanism of the CRISPR–Cas12a system specificity, we measured the melting temperature (Tm) of purified DNA/crRNA duplexes used in this study by thermal denaturation assay (41) (**Table S2**). The Tm values were slightly increased for split crRNAs compared to full ones, which may indicate that the absence of hairpin structure like scaffold domain has a positive impact on stabilization of DNA/crRNA duplexes.

Interestingly, for chimeric RNA-DNA versions bearing four deoxynucleotides in the middle (positions 13-16) of the spacer region, Tm of the duplexes was approximatively the same as for non-modified split crRNA, and only slightly decreased for the version with 8 deoxynucleotides in positions 13-20. Meanwhile, for the chimeric crRNA bearing four deoxynucleotides at the 3’ end of spacer region, split D(17-20), Tm dropped by 2 to 3°C, indicating an important decrease in duplex stability. Noteworthy, DNA:RNA duplexes normally adopt the helical conformation A, which are different from the B-form of DNA:DNA double strand helices (65). Even so, the chimeric molecules containing RNA nucleotides followed by DNA nucleotides have been found to adopt the A-form (66). We can hypothesize that the stretch of deoxynucleotides in the middle of crRNA spacer region does not change the helix conformation; in contrast, dGdAdGdT sequence at the 3’-end of the spacer has a potential to adopt the B-form, which leads to the partial melting and destabilization of the crRNA:DNA duplex.

Importantly, for non-modified and modified crRNAs, we detected the same level of Tm changes for target (cognate) and off-target (mismatched) duplexes (**Table S2**). Therefore, the Tm values, corresponding to the stability of DNA/crRNA duplexes, do not correlate with ability of crRNA-Cas12a system to distinguish between wild-type and mutant mtDNA. To explain this discrepancy, we can speculate that different regions of the spacer crRNA domain have different impact on the activity and specificity of the system, which corroborates with peculiar mechanism of target DNA cleavage by Cas12a nuclease. As we already mentioned, a single nuclease domain RuvC cleaves first the non-target strand (NTS) which is followed by target strand (TS) cleavage (**Fig. 1A**) (6,7). Molecular mechanism of the TS break is not completely understood. It was suggested that crRNA/Cas12a complex initiates template DNA cutting by forming a 20-nucleotide R-loop in which the crRNA displaces the non-target strand of a double-helical DNA substrate, positioning the RuvC nuclease active site for NTS strand cleavage (**Fig. 1B**) (7). Then, the target strand (TS) needs to bend, accessing the most probable catalytic orientation (6). Recently, detailed structural studies revealed that a helical lid of RuvC domain, after cleavage of the non-target strand DNA, adopts an open conformation and anchors the crRNA:TS duplex at PAM-distal site. Then, cooperative motions of the Cas12a domains bend the TS DNA, and the helical lid guides it toward the RuvC catalytic core for cleavage (67). It was also suggested that Cas12a exploits local duplex instability at the crRNA-3’ side of the R-loop for the strand separation of crRNA:TS duplex and target strand cut (7). This model suggests a special role of the PAM-distal part of the crRNA:DNA duplex, supporting our data on the effect of DNA substitutions in this crRNA region on the cleavage specificity (**Fig. 3**).

### Influence of magnesium concentration

The cleavage mechanism described above was also supported by smFRET approach, which directly demonstrated that the local DNA unwinding in the PAM-distal region and Mg^2+^-mediated stabilization of unwound conformation is the essential prerequisite for the target DNA cleavage reaction (59). Thus, the importance of Mg^2+^ cations for Cas12a nuclease functioning have been extended: in addition to Mg^2+^ ions playing a catalytic role (68), an unexpected structural role of Mg^2+^ ions have been revealed, namely stabilization the unwound conformation of the PAM-distal DNA and thereby promotion of the TS cleavage.

Our data show that the increase in the Mg^2+^ concentration from 1 mM to 5 mM caused 1.5-fold more efficient DNA cleavage (**Fig. 4**), which is in agreement with previously published data (58,59,69). Surprisingly, the increase in the Mg^2+^ concentration significantly affected the accuracy in DNA cleavage, as the ratio between cleavage of cognate and non-cognate DNA templates (referred to as discrimination coefficient, which is close to 1 for completely tolerated mutations) changed from 0.7-0.8 at 5 mM Mg^2+^ to 0.2-0.5 at 1 mM Mg^2+^. This discrepancy between Cas12a enzymatic activity and accuracy can be explained by the dual function of Mg^2+^ cations: catalytic role of magnesium ions coordinated in the nuclease active center and their structural role in PAM-distal region of crRNA:DNA duplex. The high Mg^2+^ concentration (5-10 mM) is optimal for catalysis but facilitates off-target cleavage due to stabilization of the mismatched crRNA:DNA duplexes, especially for DNA targets bearing mismatches in the PAM-mid and distal region. In contrast, at 1 mM Mg^2+^, the enzymatic activity is sustained at sufficient level, but the mismatches destabilizing crRNA:DNA duplex can no longer be tolerated.

This can explain the improved accuracy in DNA cleavage for Cas12a/crRNA complexes bearing chimeric crRNA D(17–20) under low Mg^2+^ concentration (**Fig. 5**). Interestingly, DNA substitutions in the middle part of crRNA D(13–16) increased mutations discrimination at 5 mM Mg^2+^ (**Fig. 3B**) but blocked the nuclease activity at low magnesium despite the same melting properties (**Table S2**). These data indicate on a local destabilization of the mid part of crRNA:DNA duplex by deoxynucleotides, which can be partially compensated by high Mg^2+^ concentration, providing higher precision of DNA cleavage. Nevertheless, at 1 mM Mg^2+^, chimeric split crRNA D(13–16) cannot provide a correct docking of crRNA/Cas12a complex on the target DNA and/or the correct position of TS DNA for its cleavage.

Our data corroborate recently published study demonstrated that CRISPR-Cas12a exhibits metal-dependent specificity switching (70). Using plasmid libraries with mismatches and phage escape assays, it was shown that lowering Mg^2+^ concentration decreased the tolerance to the mismatches in the PAM-proximal seed region, while increasing the discrimination of the PAM-distal mismatches.

In contrast to the positive effect of high Mg^2+^ concentrations on AsCas12a nuclease activity, we observed a reduced deamination activity of Cas12a-fused base editors at 5 mM MgCl_2_ compared to 1 mM MgCl_2_ (**Fig. S3D**, **Fig. S4**). This may be attributed to the particular mechanism of single-strand base editing by cytosine deaminase (61), for which the DNA duplex stabilization at higher Mg^2+^ concentrations should have a negative impact.

These results reveal the importance of physiological cations concentration for the specificity of Cas12a effector nucleases. Previous *in vitro* studies typically used 5-10 mM Mg^2+^ concentration and have shown that Cas12a is tolerant of multiple PAM-distal mutations (71). Nevertheless, the Mg^2+^ concentration is expected to be lower (∼1mM) in bacterial (57) and mammalian (72) cells and mitochondria (58) and can fluctuate, for instance, when nucleotides that coordinate Mg^2+^ are depleted (73).

In a recent study, trans-cleavage Cas12a activity have been applied to detect mtDNA mutations (58). It was revealed that non-specific ssDNA degradation (trans-cleavage) was very low at 1mM Mg^2+^; moreover, no degradation under these conditions was detected in (70). Therefore, we can speculate on more specific target DNA cleavage by Cas12a system *in vivo*, and a quasi-absence of its trans-cleavage activity, which may be activated only in case of local increase of magnesium concentration. Not only divalent cations, but also monovalent ions concentrations, fluctuations of pH, oxidation conditions and temperature may influence the Cas12a activity in different cellular compartments, which should be taken into account for the further development of the mitochondrially targeted crRNA/Cas12a system.

These observations show that the Cas12a effector specificity strongly depends on test conditions. We demonstrated that AsCas12a/crRNA system capacity to discriminate between wild type and mutant mtDNA molecules harboring pathogenic point mutations m.13513G>A and m.13514A>G can be dramatically increased by use of split chimeric crRNA bearing deoxynucleotides in the distal region of the spacer part. While these findings establish Cas12a specificity *in vitro*, an important remaining question was whether the system could also function in human mitochondria.

### *In vivo* evidence of modified crRNAs activity in human mitochondria

Recent publications demonstrated that for the mtDNA engineering approaches, linearized mtDNA should be protected by inactivation of the replisome exonuclease MGME1 (20), as the rapid degradation of cleaved mtDNA in human cells makes challenging to prove target mtDNA cleavage *in vivo*. Since the base editing approach could help to demonstrate the activity of CRISPR-Cas12a system in human mitochondria without mtDNA cleavage, we designed a version of cytosine base editor (34) based on Cas12a inactivated nuclease able to be imported into mitochondria.

First, we tested the possibility to use modified crRNAs for mtDNA base editing *in vitro*. Split and D(17–20) modified crRNAs were able to target the mtCBE base editor to the mitochondrial gene ND5 and to induce the expected base changes with efficiency only slightly lower for the split crRNA version (**Figs. 6A**, **S3D, Table S3**). Noteworthy, in cultured cells transfected with mtCBE-expressing plasmid and crRNAs, the expected base editing was significantly detectable, but its level was very low (**Fig. 6B**, **D**). The frequency of mtDNA editing in the cultured cells was low even at positions 7 and 10 of the editing window (2×10^-4^). Recently, CRISPR technology was successfully applied to edit yeast mtDNA (74). This study obtained the average editing frequency of 4×10^-9^ quantified by use of survival-based reporter. We believe that these and our data provide a proof-of-concept for the use of a CRISPR-Cas system in mitochondria.

One of the possible options to increase the editing efficiency may be the further protein engineering of base-editing enzymes. First generations of base editors relied on introduction of nicks to the strand to which crRNA anneals, otherwise base editing wasn’t efficient (75,76). That’s why for a long time Cas12a-fused base editors were not possible, since Cas12a, as we already mentioned, has single nuclease domain. However, extensive protein engineering led to a creation of highly efficient deaminases, superior to previous versions several hundred-fold (44,61). The possibility of introducing nicks to the non-deaminated strand would help with the efficiency of editing, guiding reparation systems in favor of introduced change (77).

Nevertheless, the main reason of the low editing efficiency can probably be a very small level of crRNA import into mitochondria, as it was demonstrated by our and other laboratories (17,19,74). In fact, only few percent of some non-coding cytosolic RNAs can be detected in the mitochondrial matrix (14). To address the crRNA mitochondrial delivery issue, various approaches could be tested, as, for instance, the lipophilic conjugates (78), nanoparticles (79) and more recently developed nanoflower cargos (80). Another promising approach consists in crosslink between AsCas12a effector nuclease and crRNA as described in (81). This can provide a tool for efficient crRNA delivery into human mitochondria in complex with AsCas12a protein, thus addressing the main limitation of the mitochondrial CRISPR-Cas systems.

In summary, we report that split chimeric crRNA bearing deoxynucleotides can strongly increase the capacity of the mito-AsCas12a/crRNA system to discriminate between wild-type and mutant mtDNA molecules harboring pathogenic point mutations. Furthermore, mito-AsCas12a fused to a cytosine base editor enabled targeted editing of the ND4 and ND5 genes of mtDNA in cultured cells. Further optimization? of crRNA delivery may facilitate the development of this system as therapeutic tool for shifting heteroplasmy or directly correcting pathogenic mtDNA mutations, thereby improving mitochondrial function.

## Data availability

All data presented in the manuscript are available upon request. The NGS data from this study are available in the NCBI Sequence Read Archive (SRA) under accession code PRJNA1526067.

## Acknowledgements

The authors acknowledge Anne Friedrich and Gauthier Brach (UMR7156 GMGM, CNRS/University of Strasbourg) for the help with NGS data analysis. We are grateful to Anna Smirnova (UMR7156 GMGM, CNRS/University of Strasbourg) for technical assistance in fluorescent microscopy and to Sandrine Koechler (IBMP, CNRS/University of Strasbourg) for performing NGS. This work was in the frame of the Interdisciplinary Thematic Institute IMCBio+, as part of the ITI 2021-2028 program of the University of Strasbourg, CNRS and Inserm, supported by IdEx Unistra (ANR-10-IDEX-0002), EUR (IMCBio ANR-17-EUR-0016) and SFRI (STRAT’US project, ANR-20-SFRI-0012) within the framework of France 2030 National Program. Collaboration between GMGM and ICBFM was supported by the PRC grant CNRS-RFBM mitoTARN N°297439-2019016. A.P., I. P. and M. M. were supported by the State Assignment of the Institute of Chemical Biology and Fundamental Medicine, Siberian Branch, Russian Academy of Sciences, N° <u>125012300656-5</u>. Figures were created with BioRender.com.

## Author contributions

N.S, A.M.H. conducted the experiments, N.N. contributed to crRNA design and NGS data analysis, N.S. performed experimental design, data analysis, data visualization and drafting, A.P., I. P. and M. M. performed crRNA chemical synthesis and melting temperatures analysis, N.E. and I.T. were responsible for conceptualization, data analysis, writing, reviewing and editing, I.T. and N.E. were responsible for funding.

## Competing interests

The authors declare no competing interests.

## Materials & Correspondence

Correspondence and material requests should be addressed to Nina Entelis.

## Notes

https://www.ncbi.nlm.nih.gov/bioproject/PRJNA1526067

### Competing Interest Statement

The authors have declared no competing interest.

