## Supplementary material for "Modified crRNA variants for precise targeting of point mutations in human mitochondrial DNA by CRISPR-Cas12a": Figure S1, Figure S2, Figure S3, Figure S4, Table S1, Table S2, Table S3, Table S4

### SUPPLEMENTARY INFORMATION

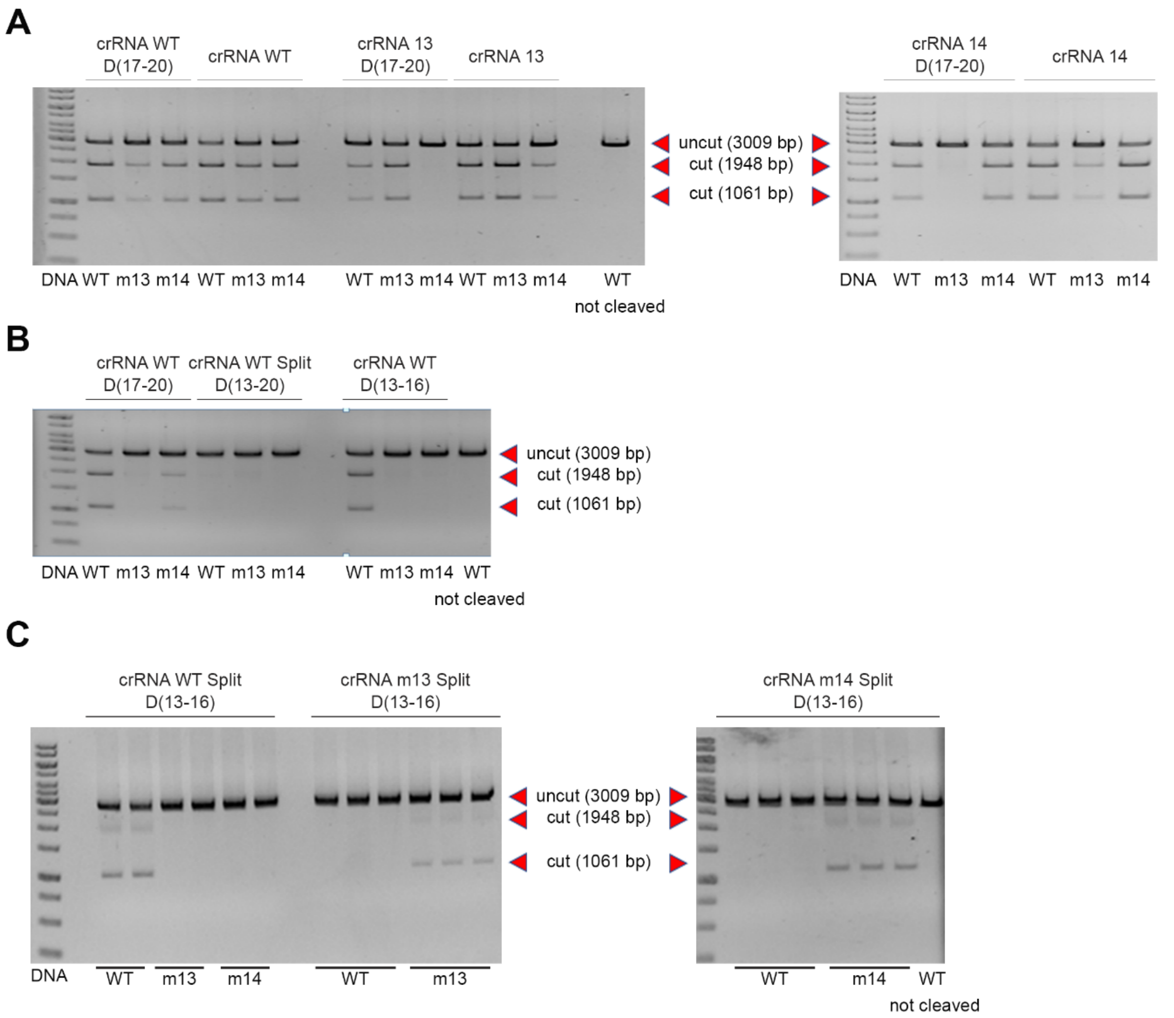

**Figure S1. Specific cleavage of mtDNA fragment by AsCas12a effector programmed with various crRNAs in buffer with 5 mM $MgCl_2$.** Representative agarose gels. **A,** *In vitro* cleavage test of target DNA (WT, m13, m14 as indicated below the panels) by AsCas12a effector programmed with non-modified and D(17-20) crRNAs indicated above the panels; n=3, representative agarose gels, DNA fragments size shown by red arrows. Corresponds to Figure 2. **B,** as in **A,** with split crRNA versions bearing deoxynucleotides in different positions, D(17-20), D(13-20) and D(13-20) (Fig 3, Table S1). **C,** cleavage of target DNA by AsCas12a with split D(13-16)WT n=2; split D(13-16)m13 and split D(13-16)m14 n=3; Fig 3B.

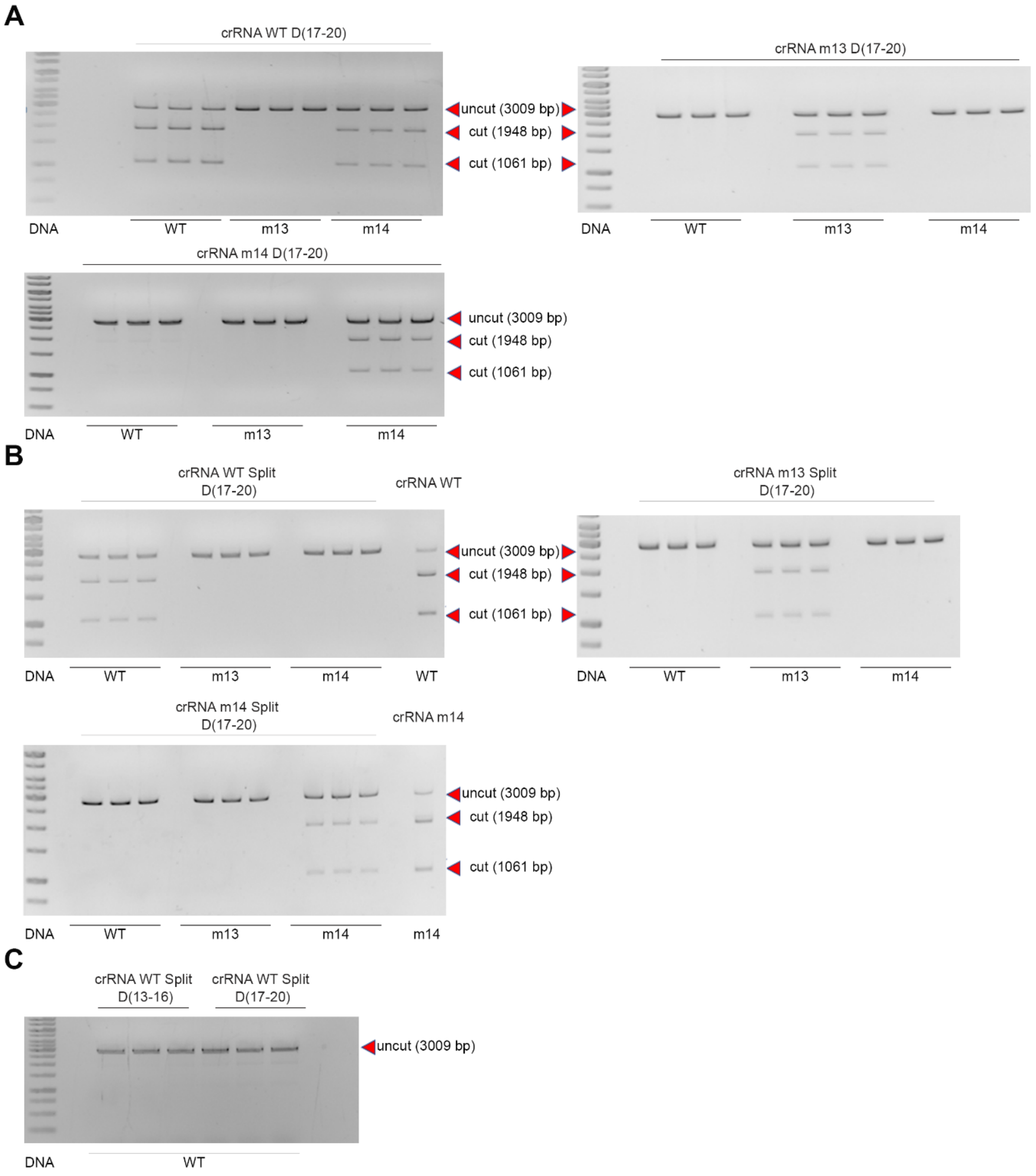

**Figure S2. Specific cleavage of mtDNA fragment by AsCas12a effector programmed with various crRNAs in buffer with 1 mM $MgCl_2$.** Representative agarose gels. **A,** cleavage of different target DNA (WT, m13, m14 as indicated below the panels) by AsCas12a programmed with crRNA D(17-20), as indicated at the left; n=3, quantification is provided in Fig. 5. **B, as in A,** with split crRNA D(17-20) as indicated above the panels. **C,** cleavage of target DNA WT with split crRNA D(13-16)WT and D(17-20)WT as indicated above the panels.

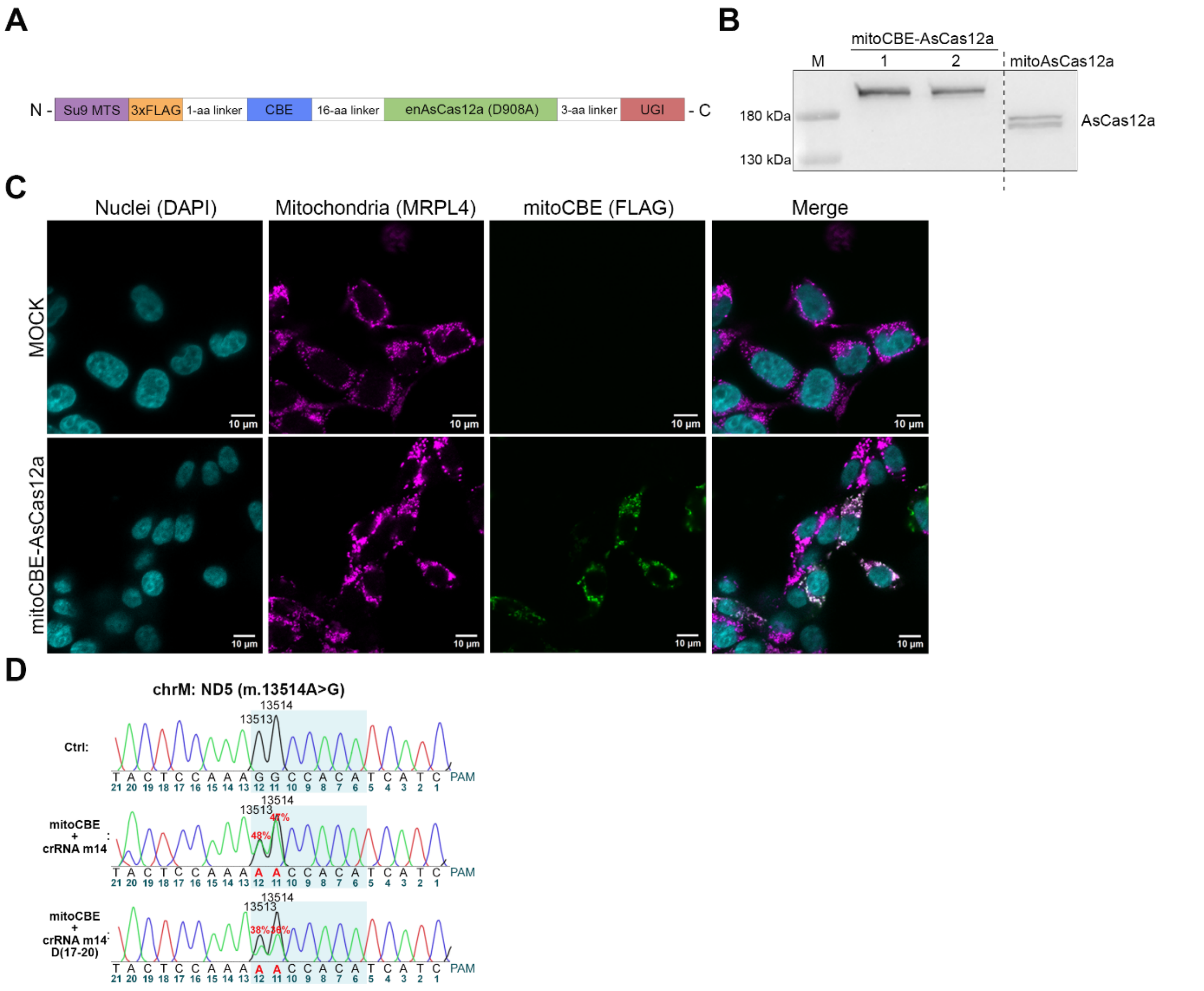

**Figure S3. Characterization of mitoCBE-AsCas12a expression and localization. A.** Schematic representation of mitoCBE-AsCas12a base editor (mtCBE) designed and used in the study. Su9 MTS, 3xFLAG epitope tag, and cytidine base editor (CBE, APOBEC3) are fused to the N-terminus of enAsCas12a (23), inactivated by a D908A mutation (dAsCas12a). Uracil-DNA glycosylase inhibitor (UGI) is attached to the C-terminal end of the base editor. Resulting construct was cloned into pET30a (+) for protein purification and into pcDNA5/FRT vector for transfection of cultured cells. **B.** Western blot analysis of HEK293 cells transfected with plasmids encoding mitoCBE-AsCas12a or mitoAsCas12a. Antibodies against AsCas12a (Table S4) were used for detection. M denotes the pre-stained protein ladder. Dashed line indicates the site of membrane cut. n=2 for mitoCBE-AsCas12a. **C.** Fluorescence microscopy of HEK293 cells expressing mtCBE. MOCK, control transfection without plasmid DNA. The 3xFLAG-tagged base editor was stained with FLAG-tag/Alexa Fluor 488 antibodies (green), mitochondria were stained with MRPL4/Alexa Fluor 647 antibodies (magenta), and nuclei were stained with DAPI (blue). Scale bars: 10 $\mu$m. **D.** Sanger sequencing analysis of a PCR-amplified HEK293 mtDNA fragment treated *in vitro* with mitoCBE-AsCas12a in complex with various crRNAs m14 as indicated at the left, in buffer containing 1mM $Mg^{2}$. The percentage of mutations introduced by the base editor is shown in red. The mitoCBE-AsCas12a editing window is highlighted in blue. An untreated PCR template served as the control. n=1.

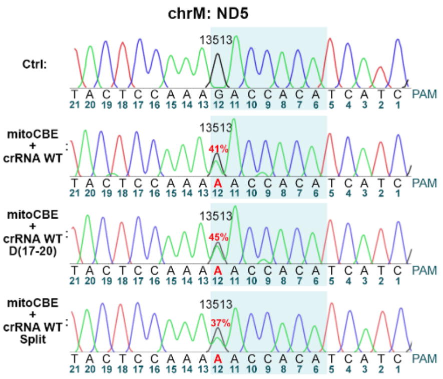

**Figure S4: Application of mitoCBE-AsCas12a to the MT-ND5 gene *in vitro*.** Sanger sequencing analysis of a PCR-amplified HEK293 mtDNA fragment treated *in vitro* with mitoCBE-AsCas12a in complex with crRNAs in buffer containing 5mM $Mg^{2+}$. Unmodified crRNA WT, crRNA containing deoxynucleotides at positions 17–20 (D17–20) and a split crRNA (indicated at the left) were used to introduce the m.13513A>G mutation in ND5 mtDNA gene. The percentage of mutations introduced by the base editor is indicated in red. The mitoCBE-AsCas12a editing window is highlighted in blue. An untreated PCR template served as the control. n=2.

**Table S1. Sequence of crRNAs and DNA oligonucleotides.**

The crRNA spacer sequences are shown in green; deoxynucleotides are colored in blue. The scaffold sequence is underlined. The positions of m13 and m14 mutations are highlighted in bold.

| Name | Sequence 5’-3’ |
|---|---|
| | **crRNA** |
| **WT** | UAAUUUCUACUCUUGUAGAUGAUGAUGUGGUCUUUGGAGU |
| **m13** | UAAUUUCUACUCUUGUAGAUGAUGAUGUGGU**U**UUUGGAGU |
| **m14** | UAAUUUCUACUCUUGUAGAUGAUGAUGUGG**C**CUUUGGAGU |
| **WT D(17-20)** | UAAUUUCUACUCUUGUAGAUGAUGAUGUGGUCUUUGdGdAdGdT |
| **m13 D(17-20)** | UAAUUUCUACUCUUGUAGAUGAUGAUGUGGU**U**UUUGdGdAdGdT |
| **m14(17-20)** | UAAUUUCUACUCUUGUAGAUGAUGAUGUGG**C**CUUUGdGdAdGdT |

| | |
|---|---|
| **Spacer WT** | GAUGAUGUGGUCUUUGGAGU |
| **Spacer m13** | GAUGAUGUGGU**U**UUUGGAGU |
| **Spacer m14** | GAUGAUGUGG**C**CUUUGGAGU |
| **Scaffold** | UAAUUUCUACUCUUGUAGAU |
| **Spacer WT D(17-20)** | GAUGAUGUGGUCUUUGdGdAdGdT |
| **Spacer m13 D(17-20)** | GAUGAUGUGGU**U**UUUGdGdAdGdT |
| **Spacer m14 D(17-20)** | GAUGAUGUGG**C**CUUUGdGdAdGdT |
| **Spacer WT D(13-16)** | GAUGAUGUGGUCdTdTdTdGGAGU |
| **Spacer m13 D(13-16)** | GAUGAUGUGGU**U**dTdTdTdGGAGU |
| **Spacer m14 D(13-16)** | GAUGAUGUGG**C**CdUdUdUdGGAGU |
| **Spacer WT D(13-20)** | GAUGAUGUGGUCdTdTdTdGdGdAdGdT |
| **DNA primers** | |
| **EcoRI-ND5** | CCAGTGAATTCTTATGTGCTCCGGGTCCATCAT |
| **HindIII-ND5** | CGCCAAGCTTTAGGCTTCCGGCTGCCAG |
| **ND5 mut13F** | GATGATGTGG**C**CTTTGGAGTAG |
| **ND5 mut14F** | GATGATGTGGT**T**TTTGGAGTAG |
| **ND5 mut R** | GAAACCGCAAACATATCATAC |
| **DNA strands for melting experiments** | |
| **WT** | ACTCCAAAGACCACATCATC |
| **m13** | ACTCCAAA**A**ACCACATCATC |
| **m14** | ACTCCAAAG**G**CCACATCATC |

**Table S2. Melting temperatures (Tm,°C) of the crRNA:DNA duplexes.**
The experimental error of Tm is ±0.2°C.
Tm of cognate duplexes (without mismatches) are highlighted in green.

| crRNA /DNA | WT | m13 | M14 |
|---|---|---|---|
| Full-WT | 61.0 | 53.0 | 58.5 |
| Full-m13 | 54.5 | 58.0 | 51.0 |
| Full-m14 | 56 | 49.0 | 65.0 |
| Split-WT | 62.3 | 53.1 | 59.7 |
| Split-D(17-20)-WT | 59.3 | 50.0 | 56.3 |
| Split-D(17-20)-m13 | 51.5 | 55.6 | 48.6 |
| Split-D(17-20)-m14 | 54.2 | 45.6 | 62.5 |
| Split-D(13-16)-WT | 62.7 | nd | 58.9 |
| Split-D(13-16)-m13 | 55.9 | 58.8 | 52.6 |
| Split-D(13-16)-m14 | 56.7 | nd | 65.9 |
| Split-D(13-20)-WT | 61.4 | nd | 58.6 |
| Split-D(13-20)-m13 | 54.2 | 58.2 | 51.3 |
| Split-D(13-20)-m14 | 55.6 | nd | 64.7 |

**Table S3.** Results of ND5 *in vitro* deamination tests analyzed by R-based software EditR.

| $MgCl_2$ | RNA | Editing efficiency, % |
|---|---|---|
| 1mM | crRNA | 71;74 |
| | crRNA D(17-20) | 71;70;73 |
| | Split | 49;49;61 |
| | Split D(17-20) | 13 |
| 5 mM | crRNA | 41;30 |
| | crRNA D(17-20) | 46;45 |
| | Split | 37;19 |
| | Split D(17-20) | 13 |

**Table S4. Antibodies used in the study**

| Antibody's name | Provider | Reference |
|---|---|---|
| Primary antibodies | | |

| | | |
|---|---|---|
| Mouse monoclonal anti-FLAG (clone M2) | Sigma-Aldrich | F1804 |
| Rabbit polyclonal anti-MRPL4 | Proteintech Group | 27484-1-AP |
| Mouse monoclonal anti-AsCas12a | Sigma-Aldrich | MABE1823 |
| Goat polyclonal anti-Actin (C-11) | Santa Cruz Biotechnology | sc-1615 |
| Secondary antibodies | | |
| Rabbit polyclonal anti-Goat IgG HRP | Sigma-Aldrich | A8919 |
| Rabbit polyclonal anti-Mouse IgG HRP | Agilent | P0260 |
| Goat polyclonal anti-Mouse IgG-Alexa Fluor 488 | Thermo Fisher Scientific | A-11001 |
| Goat polyclonal anti-Rabbit IgG-Alexa Fluor 647 | Thermo Fisher Scientific | A-21246 |
